# In-Cell Sensitivity-Enhanced NMR of Intact Living Mammalian Cells

**DOI:** 10.1101/2021.05.28.446194

**Authors:** Rupam Ghosh, Yiling Xiao, Jaka Kragelj, Kendra K. Frederick

## Abstract

NMR has the resolution and specificity to determine atomic-level protein structures of isotopically-labeled proteins in complex environments and, with the sensitivity gains conferred by dynamic nuclear polarization (DNP), NMR has the sensitivity to detect proteins at their endogenous concentrations. However, DNP sensitivity enhancements are critically dependent on experimental conditions and sample composition. While some of these conditions are theoretically compatible with cellular viability, the effects of others on cellular sample integrity are unknown. Uncertainty about the integrity of cellular samples limits the utility of experimental outputs. Using several measures, we establish conditions that support DNP enhancements that can enable detection of micromolar concentrations of proteins in experimentally tractable times that are compatible with cellular viability. Taken together, we establish DNP assisted MAS NMR as a technique for structural investigations of biomolecules in intact viable cells that can be phenotyped both before and after NMR experiments.

**Classification:** Biophysics and Structural Biology

## Introduction

Despite the importance of the environment, structural investigations of biomolecules are typically confined to highly purified *in vitro* systems. These investigations yield invaluable insights, but such investigations cannot fully recapitulate cellular environments. Capturing the effect of these complicated environments on biomolecular conformation is of particular importance for proteins that have more than one stable conformation, interact with cellular components or contain regions of intrinsic disorder.^1^

In-cell NMR allows us to obtain atomic resolution information of proteins in their native environments.^2-8^ Nuclear Magnetic Resonance (NMR) spectroscopy detects only NMR-active nuclei. These nuclei are non-perturbative, label-free probes that can be specifically incorporated into a protein of interest that is either delivered to or expressed inside the cell.^5-9^ In-cell NMR has been employed to investigate protein and nucleic acid structure, dynamics, interactions, and post-translational modifications like phosphorylation.^4,8-21^ Solution state NMR spectroscopy can be used to study interactions in cells however, this technique is limited to small proteins that interact with small ligands or only transiently with larger biomolecules. This is because solution state NMR spectroscopy is limited by molecular tumbling times that depend upon molecular size and solvent viscosity and thus reports only on rapidly re-orienting molecules in the sample.

Magic angle spinning (MAS) solid-state NMR is not limited by molecular correlation times and is particularly useful to study large protein complexes, amyloid fibrils and membrane proteins.^21-26^ With the sensitivity gains conferred by dynamic nuclear polarization (DNP), MAS NMR has the sensitivity to detect proteins at their endogenous concentrations in complex biological environments.^1,21,22,27^ However, the effectiveness of DNP-enhanced MAS NMR is critically dependent on sample composition^28-34^ and experimental conditions^23,35-38^. DNP increases the sensitivity of NMR spectroscopy through the transfer of the large spin polarization of an unpaired electron to nearby nuclei^35^ which are typically introduced into a sample by doping with millimolar concentrations of stable biological radicals^32-34^. In addition to millimolar concentrations of polarizing agents, a typical DNP sample of a hydrated biomolecule is cryoprotected by the addition of 60% *d*_*8*_-glycerol to aid in the formation of vitreous ice, deuterated to ∼90% to aid spin diffusion, frozen to near liquid nitrogen temperatures and subjected to magic angle spinning. Pioneering work applying DNP MAS NMR to cultured mammalian cells suggested that measurement of low concentrations of proteins inside mammalian cells is possible^22^ but uncertainty about the biological integrity of the cellular sample limits the utility of the structural information.

Here, we establish an approach to homogenously introduce the polarization agent, AMUPol^32^, into cultured mammalian cells that results in high sensitivity gains and supports cellular viability throughout DNP NMR experimentation. After structural characterization via DNP MAS NMR, these cells can be cultured or imaged and their phenotype can be determined and compared with cells before structural characterization. Our work sets the stage to determine if and how cellular environments modulate protein structure, which is of particular interest for systems where protein conformation can affect cellular fate. Investigation of protein conformations inside viable cells using DNP MAS NMR creates an experimental system with the ability to tightly couple genotypes, phenotypes and environments (e.g. presence/absence of a drug) to specific structures or structural ensembles.

## Results

### HEK293 cells remain viable during DNP MAS NMR

#### Conditions that favor efficient DNP and HEK293 cell viability are compatible

To determine whether any of the components that are typically used to achieve high DNP-enhancements for biological systems compromise cellular viability, we used a trypan blue dye exclusion test to determine the percentage of cells with intact membranes present in a sample. Unsurprisingly, HEK293 viability was not compromised by replacement of media components with PBS, by addition of common working concentrations of cryoprotective agents (10% DMSO or 15% glycerol)^39^, or by either per-deuteration or complete deuteration of the buffer components (Figure S1A-C). Moreover, HEK293 viability was not compromised by addition of the polarization agent AMUPol at concentrations up to 50 mM (Figure S1E) even after 45 minutes of exposure at room temperature (Figure S1F). To ensure that AMUPol, or any of the other treatments, were not toxic to HEK293 cells, we assessed viability using an orthogonal method; a quantitative cell growth assay. HEK293 cells exposed to glycerol had a lag phase before exponential growth however the growth kinetics of the exponential phase were indistinguishable from untreated HEK293 cells once exponential growth began (Figure 1C).

**Figure 1:**
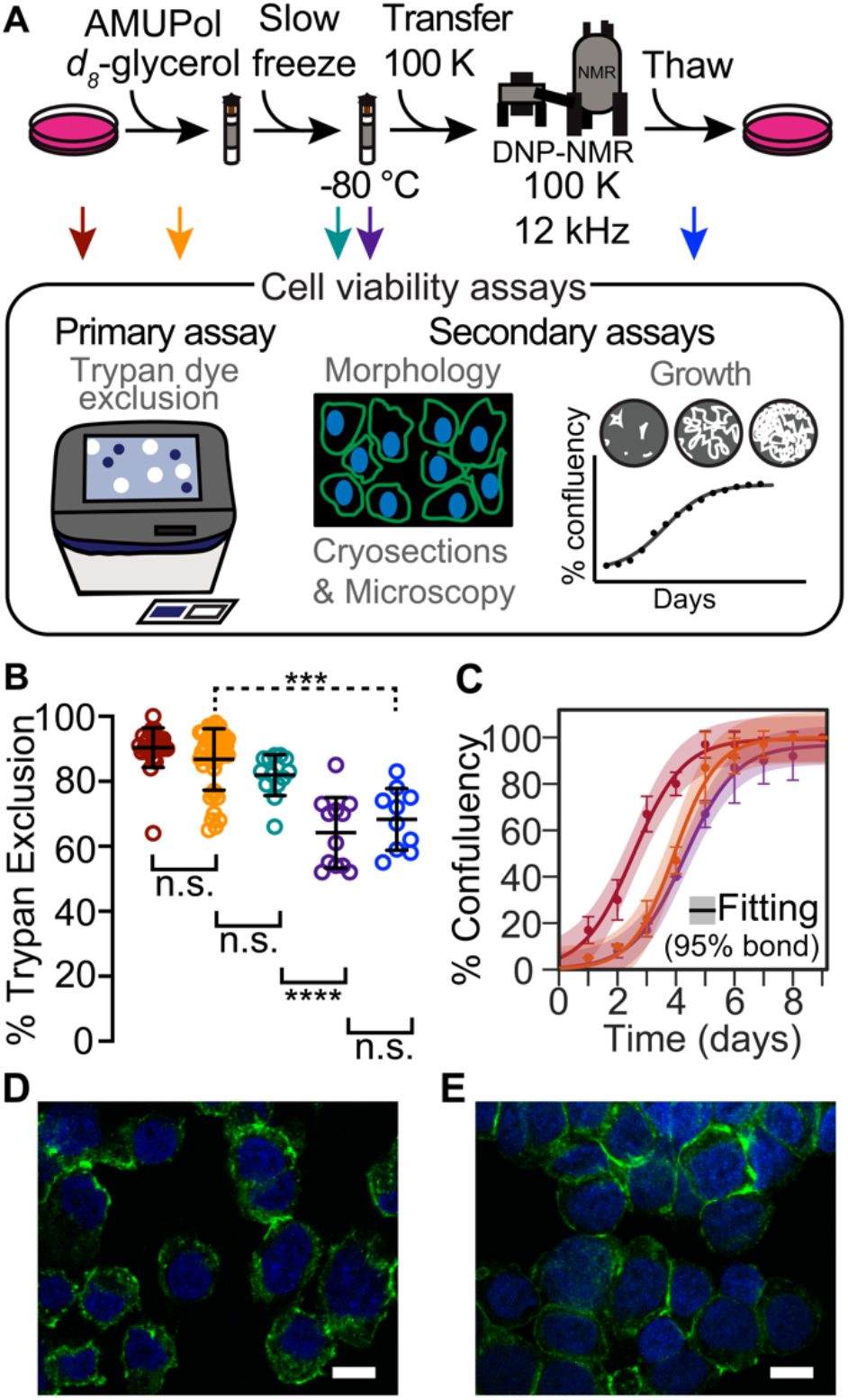
HEK293 cells are viable throughout the DNP NMR process. (A). Experimental scheme of the DNP NMR sample preparation procedure and cell viability assays. Colored arrows indicate points at which sample viability was assessed. Viability was assessed for cells after trypsinization and washing (dark red), after suspension in AMUPol and cryoprotectants (orange), after being from at 1 °C per min (green), after manipulation to remove the drive tip from the frozen rotor (purple) and after the entire DNP MAS NMR experiment (blue). (B) Frozen cells best represent the state of the sample during DNP NMR data collection. Percentage of cells with trypan impermeable membranes at each sample assessment point, colored as in A. Each point represents an independent sample. Black bars indicate average and standard deviation. Brackets indicate results of two-tailed homoscedastic student’s t-tests. Unpaired comparisons are indicated with solid brackets and paired comparisons with dashed brackets. (n.s. *p* > 0.05, * *p* < 0.05, *** *p* < 0.001, **** *p* < 0.0001). (C) Addition of glycerol affects growth kinetics, but further sample manipulations do not. Growth kinetics as assessed by confluency, colored as in A. The averages and standard deviations of three independent experiments are indicated by circles and error bars, respectively. The best fit of sigmoid is indicated in solid lines and the 95% confidence interval by the shaded area. (D and E) Fluorescence microscopy of HEK293 cells before (D) and after freezing (E) using Phalloidin (green) and Hoechst 33342 (blue) to stain actin cytoskeleton and nucleus (Scale bar 10 µm).

Neither per-deuteration nor exposure to AMUPol had any additional effects on cell growth (data not shown). Thus, HEK293 cell viability is not compromised by the components that support high DNP-enhancements. However, the amount of added cryoprotectants can affect viability; suspension of cells in 60% glycerol, a percentage commonly used in biological DNP samples, resulted in the loss of membrane integrity of more than half of the cells (Figure S1C). To determine whether any of these components compromise cellular viability during or after cryopreservation, we used a trypan blue dye exclusion test to determine the percentage of viable cells after freezing in the presence of these compounds. Unsurprisingly, freezing cells that were cryoprotected in DNP-compatible media by direct immersion of the sample in liquid nitrogen (“flash freezing”) resulted in fewer than 20% of cells with trypan blue impermeable membranes, none of which were able to grow when the sample was plated (Figure S1D). However, freezing cells that were cryoprotected in DNP-compatible media at the standard controlled rate of 1 °C/min had high post thaw viabilities (Figure 1E, Figure S1D) and had normal growth kinetics after being thawed (Figure 1C). Thus, HEK293 cells can be cryopreserved in matrices that support high DNP-enhancements in biological samples.

#### Cryogenic transfer of slowly frozen cellular samples supports post-experiment viability

Because freezing rate affects the success of cryopreservation, we next assessed various methods of transferring cellular samples into the NMR spectrometer. We first assessed two common sample handling methods for low temperature MAS NMR; transfer of a room temperature sample into a pre-cooled instrument and cryogenic transfer of a flash frozen sample into a pre-cooled instrument^40^. Transfer of a room temperature rotor into an NMR probe that is pre-equilibrated to 100 K reduces the sample temperature by ∼200 K in ∼2 minutes followed by a slow decrease to 100 K over 15 minutes. Freezing the sample inside of the spectrometer in this manner resulted in fewer than 5% viable cells at the end of the experiment. Flash freezing of the sample by immersion of the room temperature rotor into a liquid nitrogen bath followed by a cryogenic transfer into spectrometer^37^ resulted in fewer than 20% viable cells at the end of the experiment. As expected, freezing rates on these timescales resulted in substantial loss of cellular viability^40,41^. Thus, we froze cryoprotected cells at the controlled rate of 1 °C/min and then cryogenically transferred the sample from liquid nitrogen storage directly into a spectrometer that was pre-equilibrated to 100 K.^40^ Cryogenic transfer of slowly frozen sample resulted in high post DNP viability (Figure 1B).

#### Neither MAS nor DNP affect cellular viability

To determine whether the any of the manipulations required for DNP MAS NMR sample preparation compromise cellular viability, we assessed trypan blue membrane permeability at each step of our sample preparation workflow (Figure 1A, arrows). After harvesting adherent cells from tissue culture plates, the cells were rinsed with PBS and pelleted. At this point, cellular membrane integrity as assessed by trypan blue dye exclusion tests was high (92 ± 3%, Figure 1, dark red). Addition of cryoprotectants and AMUPol followed by transfer into 3.2 mm NMR rotors slightly reduced membrane integrity (87 ± 9%, Figure 1, orange; decrease of 5 ± 10%, *p* = 0.03). Freezing cryoprotected cells at the controlled rate of 1 °C/min did not significantly compromise membrane integrity as assessed by trypan blue dye exclusion test (82 ± 6%, Figure 1, green, *p* = 0.09). Post-NMR, trypan blue membrane integrity was lower than those for slow frozen samples (68 ± 9%, Figure 1, blue; decrease of 14 ± 11%, *p* = 4e-4). Interestingly, however, when membrane integrity was determined both pre-freezing (Figure 1A, orange arrow) and post-NMR experiment (Figure 1A, blue arrow) on the same sample – as opposed to comparing the averages - the decrease in viability was smaller (decrease of 11 ± 3%, *p* = 0.001, *n* = 10, rather than 19 ± 13%). This suggested that sample specific differences in rotor packing, DNP MAS NMR conditions, or unpacking may affect the membrane integrity. To determine if the membrane integrity decrease post-NMR experiment was a result of DNP MAS NMR or sample handling, we measured membrane integrity for samples that were frozen in NMR rotors with a cap and plug, but not subjected to NMR experimentation. We found that membrane integrities of capped samples that were not subjected to NMR were indistinguishable from samples that were subjected to NMR experimentation (64 ± 11%. Figure 1, purple, *p* = 0.35). Thus, the post-NMR decrease in membrane integrity was not a result of DNP MAS NMR but rather was a result of the sample handling required to remove the drive tip and silicon plug before the membrane integrity could be measured. Thus, cryoprotected frozen samples, rather than post-NMR samples, are most representative of the state of the sample when it is measured by NMR.

#### Cellular samples can be phenotyped before and after DNP MAS NMR analysis

Because membrane integrity only reports on one aspect of cellular sample integrity, we next assessed cellular integrity by two additional methods. To determine whether the any of the manipulations required for DNP MAS NMR sample preparation compromise cellular propagative ability, we next assessed cellular growth kinetics at each step in our workflow and again found that exposure to glycerol introduced a lag phase of 1.5 ± 0.5 days in growth, however no other perturbation significantly altered growth kinetics (Figure 1C). The disconnect between the membrane permeability and propagation indicated that cells with compromised membrane integrity are not necessarily dead. Finally, to determine if cells retained their morphology under these conditions, we used fluorescence microscopy to image cells before (Figure 1D and S2) and after (Figure 1E and S2) the freezing process. We observed round and intact cells with well-defined nuclei in both conditions. Taken together, this work indicated that majority of cells in cryopreserved samples for DNP were not only viable during DNP MAS NMR data collection but able to propagate after DNP MAS NMR data collection. Moreover, cellular samples can be imaged by fluorescence microscopy both before and after structural analysis by DNP MAS NMR.

### Addition of AMUPol to HEK293 cells results in DNP enhancement of all biomass components

#### Quantitative reporters of cellular biomass components

In order to assess whether the polarization agent, AMUPol, is able to increase the sensitivity of DNP experiments in intact cells, we first determined which peaks in the NMR spectra could be used as reporters of the different cellular biomass components. The majority of cellular biomass is comprised of proteins, lipids, and nucleotides (DNA and RNA). Each of these biomolecules has chemical moieties with characteristic chemical shifts (Figure 2A). Thus, as a first step, we determined whether the integrated intensity at the chemical shift of each characteristic moiety for the major biomass component could be used as a quantitative indicator of the major biomass constituents of HEK293 cells. To do so, we collected ^13^C direct polarization (DP) spectra and integrated the characteristic ^13^C peak intensity of each major biomass component to estimate their approximate quantities on samples that did not contain any polarization agent and were not subject to microwave irradiation. We found that the biomass compositions determined by ^13^C DP at 100 K for HEK293 cells were similar to the biomass composition for HEK293 cells determined by chemical means^42^ (Figure 2C and D). Thus, the integrated intensity of these characteristic chemical shifts reported quantitatively on the major biomass constituents of HEK293 cells.

**Figure 2:**
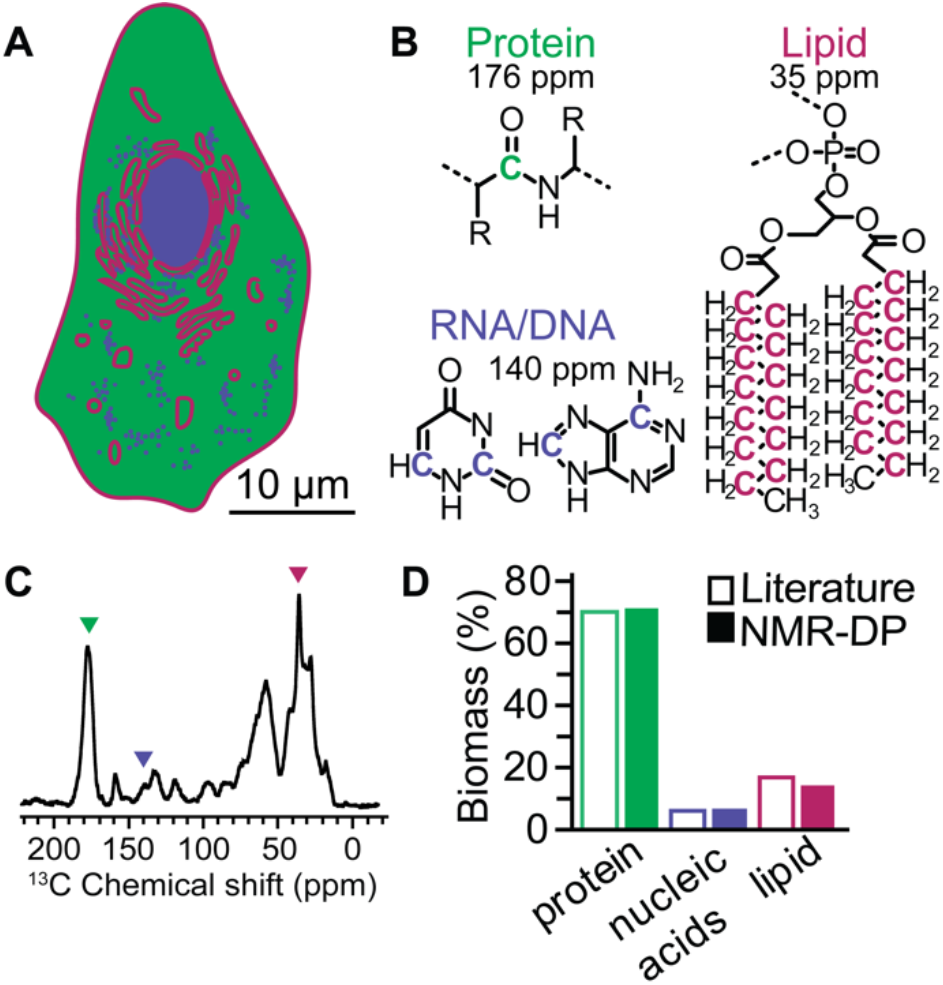
NMR report quantitively on cellular biomass composition. (A) Schematic of a mammalian cell depicting the heterogenous distribution of the three major components of the cellular biomass: protein (green), nucleotides (blue) and lipids (pink). (B) Structures of the biomass moieties with the representative carbon for each biomass component highlighted. (C) ^13^C direct polarization spectra of cryopreserved HEK293 cells grown on isotopically enriched media at 100 K taken at 600 MHz with 12 kHz magic angle spinning and a recycle delay of 300 seconds. Colored arrowheads indicate integrated peaks. (D) Quantification of biomass by NMR (closed bars) is similar to quantification of biomass by orthogonal methods^42^ (open bars).

#### DNP is very efficient for all biomass components in cell lysates

After having confirmed that the integrated intensity of the selected peaks reported quantitatively on the major biomass constituents of HEK293 cells, we next determined the DNP performance, the DNP enhancement and DNP build-up times (*T*_1_), we could expect for complex mixtures of these biomolecules. We collected ^13^C cross-polarization (CP) spectra with and without microwave irradiation, to report DNP enhancement, for lysed cells prepared with a range of AMUPol concentrations. We found that the DNP enhancements for protein in lysed cells increased with increasing concentrations of AMUPol to a maximum value of ∼93 at 5 mM AMUPol and then decreased with increasing AMUPol concentrations (Figure 3A). Taken together, these experiments on lysed cells demonstrate that AMUPol effectively enhanced all of the major biomass components.

**Figure 3:**
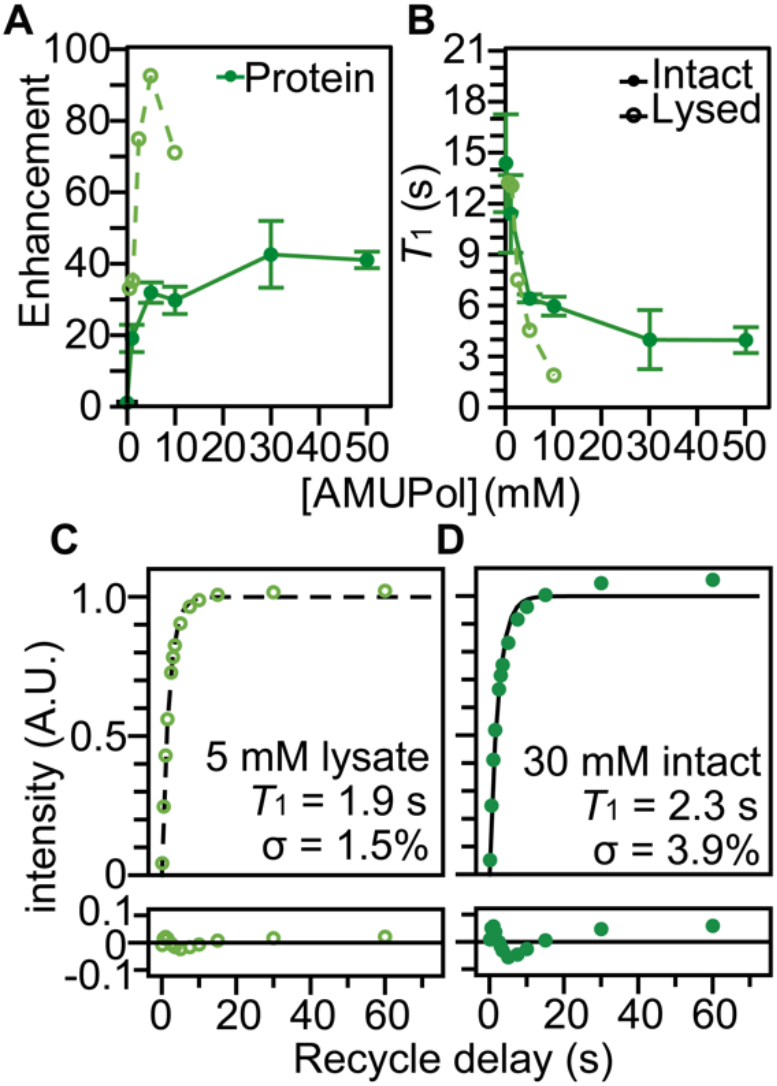
The polarization agent, AMUPol, effectively polarizes proteins in both lysed (dashed lines, light green open circles) and intact (solid lines, closed dark green circles) HEK293 cells. (A) DNP enhancement and (B) *T*1 values from saturation recovery experiments are dependent upon the AMUPol concentration. Points indicate average and error bars indicate standard deviation of the measurements. (C) Fit of the *T*1 data (open circles) for lysed cells containing 5 mM AMUPol to a mono-exponential equation (black dashed line) had a *T*1 value of 1.9 seconds with a regression error (lower plot) of 1.5%. (D) Fit of the *T*1 data (closed circles) for intact cells incubated with 30 mM AMUPol to a mono-exponential equation (black solid line) had a *T*1 value of 2.3 seconds with a regression error (lower plot) of 3.9%. Analyses for additional biomass components are presented in Supplemental Figure 3.

#### Intact viable cells have moderate DNP efficiency when incubated the AMUPol

Mammalian cells are enclosed by selectively permeable membrane that regulates the movement of molecules across the cell and they exclude many exogenous molecules outside the plasma membrane. To determine if AMUPol, a small molecule with a molecular weight of 750 Da, could effectively enhance the signal for all the components of cellular biomass for intact HEK293 cells, we measured DNP enhancements and *T*_1_ values of the major biomass components (Figure S3). DNP build-up time (*T*_1_) of NMR signal is shorter when radical concentrations increase. We collected DNP enhancements and *T*_1_ build-up times for intact cells prepared by adding increasing concentrations of AMUPol to the sample. Interestingly, we found that the DNP enhancements for proteins in intact cells reached a maximum value of 43 ± 9, which is half of the maximal enhancement attained for proteins in lysates and that the addition of 30 mM AMUPol, rather than 5 mM AMUPol, was required to attain this enhancement. The lower enhancements and longer *T*_1_ for the biomass components in intact cells relative to lysed cells indicate that the AMUPol concentration inside of the cell is lower than the sample concentration, likely because the majority of the AMUPol is excluded by the semi-permeable plasma membrane. This suggested that AMUPol is heterogeneously distributed in samples of intact cells.

#### AMUPol is heterogeneously distributed in cells incubated with AMUPol

To assess the homogeneity of the AMUPol concentration throughout each biomass component (Figure S3), we used the regression error of the fit of the *T*_1_ data to a mono-exponential equation. If the concentration distribution of AMUPol is heterogenous, there will be a mixture of underlying *T*_1_ values which will increase the regression error. The regression error of the fit of the *T*_1_ data to a mono-exponential function of the amino acid proline suspended in a matrix of 60:30:10 (v/v) glycerol:D_2_O:H_2_O with 10 mM AMUPol was 0.5% and represents the error expected from experimental noise. For lysed biomass, the regression error for protein was 1.3 ± 0.2%, for nucleotide was 1.0 ± 0.2% (lower than protein, *p* < 0.01, *n* = 6), and for lipid was 1.1 ± 0.3% (indistinguishable from protein, *p* = 0.86, *n* = 6). The increase in regression error for the lysed cells relative to that for proline reflect the larger range of underlying *T*_1_ values of the constituents in this complex mixture. For intact cells, the regression error for protein was 3.0 ± 0.8%, for nucleotide was 3.0 ± 0.8% (indistinguishable from protein, *p* = 0.62, *n* = 12), while the regression error for lipid was 2.5 ± 0.8% (somewhat lower than protein and nucleotide, *p* < 0.005, *n* = 12). The larger regression errors for the intact cells indicated that the concentration distribution of AMUPol is more heterogeneous in intact cells than in lysed cells. Taken together, this indicates that while AMUPol can polarize all of the biomass components in intact cells, the AMUPol concentration throughout the sample is heterogenous. This is likely because much of the AMUPol cannot efficiently access the interior of the cell.

### Improved delivery of AMUPol to HEK293 cells via electroporation

#### High viability of cells electroporated in the presence of AMUPol

Because only a small portion of the exogenously added AMUPol appears to be able to enter the cell, we sought to improve AMUPol delivery by transiently permeabilizing the plasma membrane via electroporation. We assessed trypan blue membrane permeability at each step in the protocol (arrows, Figure 4A) to determine whether the any of the steps during the DNP MAS NMR sample preparation of electroporated HEK293 cells compromises cellular viability. After harvesting adherent cells from tissue culture plates, the cells were rinsed with PBS, suspended in electroporation buffer and electroporated. Electroporation reduced trypan blue membrane integrity to 81 ± 9% (Figure 4, red; a decrease of 9 ± 10%, *p* = 7e-5). Electroporation in the presence of AMUPol and the addition of cryoprotectants followed by transfer into the NMR rotors did not result in any further loss in trypan blue membrane integrity 79 ± 11% (Figure 4, orange; *p* = 0.32). Freezing electroporated and cryoprotected cells at the controlled rate of 1 °C/min reduced trypan blue membrane integrity to 60 ± 15% (Figure 4, green; a decrease of 19 ± 19%, *p* = 3e-7). However, the membrane integrity after rotor unpacking and DNP MAS NMR (Figure 4, purple and blue, respectively) were indistinguishable from those for slow frozen samples.

**Figure 4:**
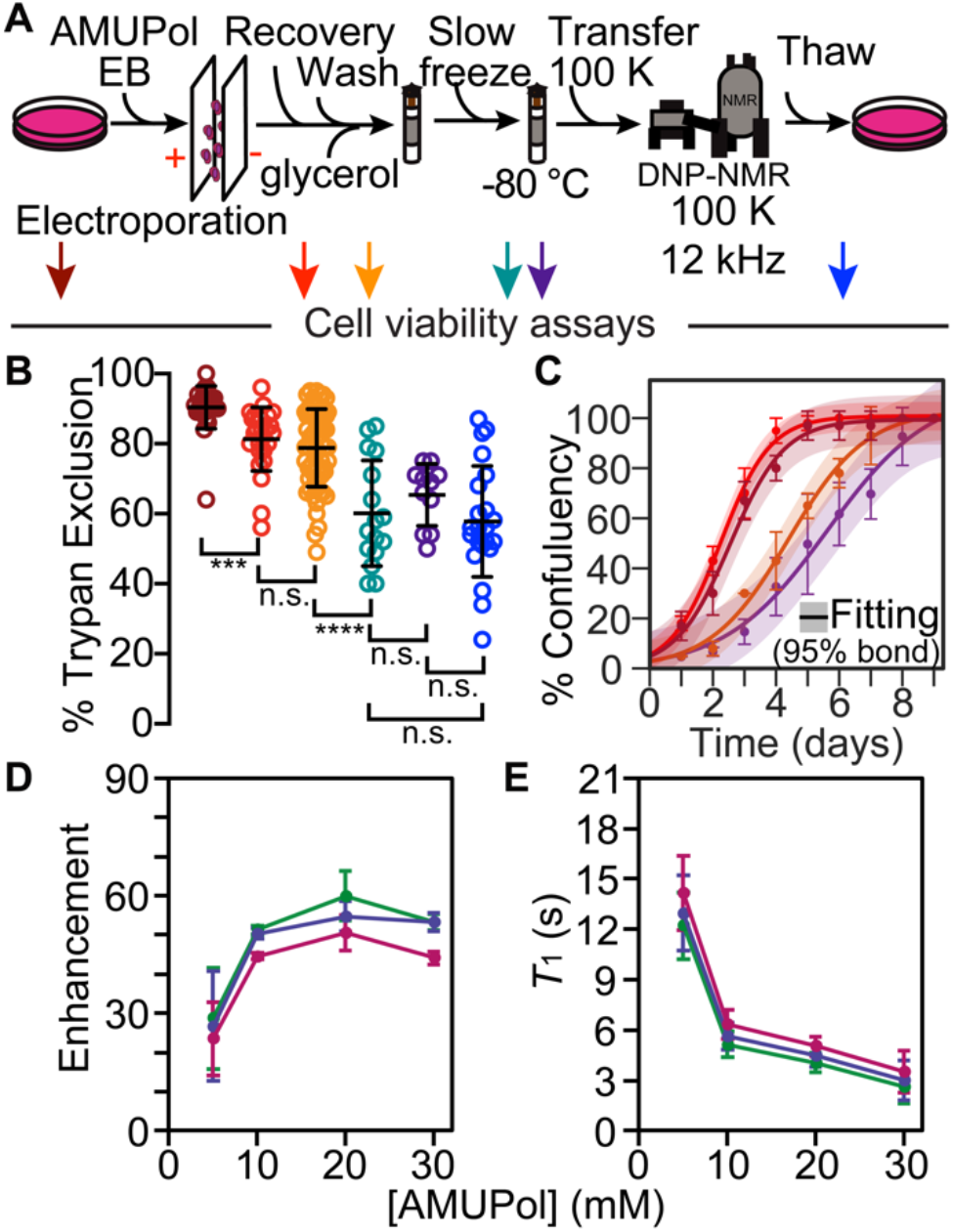
Electroporation HEK293 cells remain viable after throughout the DNP NMR process. (A) Experimental scheme of the DNP NMR sample preparation procedure that includes an electroporation step to introduce AMUPol into the sample. Colored arrows indicate points at which sample viability were assessed. Viability was assessed for cells after trypsinization and washing (dark red), after electroporation (red), after suspension in AMUPol and cryoprotectants (orange), upon thaw (green), upon thaw after manipulation to remove the drive tip from the rotor (purple) and upon thaw after DNP MAS NMR (blue). (B) Frozen cells best represent the state of the sample during DNP NMR data collection. Percentage of cells with trypan blue impermeable membranes at each sample assessment point, colored as in A. Each point represents an independent experiment. Black bars indicate average value and standard deviations. Brackets indicate comparisons for unpaired two-tailed homoscedastic student’s t-tests (n.s. *p* > 0.05, **** *p* < 0.0001). (C) Addition of glycerol affects growth kinetics, but other sample manipulations, including electroporation, do not. Growth kinetics as assessed by confluency, colored as in A. The averages and standard deviations of three independent experiments are indicated by circles and error bars, respectively. The best fit a sigmoid is indicated in solid lines and the 95% confidence interval by the shaded area. (D) DNP enhancement and (E) average *T*1 values from saturation recovery experiments for all biomass components, protein (green), nucleotide (blue) and lipid (pink), are dependent upon the AMUPol concentration at the time of electroporation; AMUPol concentration at the time of experiment will be lower. Error bars indicate standard deviations of the measurements.

#### Electroporated cellular samples can be phenotyped before and after DNP MAS NMR analysis

We next assessed growth kinetics and again found that exposure to glycerol introduced a lag phase of 2.2 ± 0.5 days (*p* = 0.02, *n* = 3) but did not affect growth rate (Figure 4C). Freezing cells increased the lag phase by 1.2 ± 0.9 days although this change not significant (*p* = 0.13, *n* = 3). However, no perturbation altered the growth rate, including electroporation (*p* = 0.7, *n* = 3) and freezing (*p* = 0.23, *n* = 3). Given that we found that membrane integrity and propagative ability were not tightly coupled for cells incubated with AMUPol, this is perhaps not particularly surprising. Taken together, this work indicated that majority of cells in a sample that is cryopreserved after delivery of AMUPol via electroporation were not only viable during DNP MAS NMR data collection but also able to propagate after DNP MAS NMR data collection.

#### Improved DNP efficiency and homogeneity when AMUPol is delivered via electroporation

To determine if introduction of AMUPol into HEK293 cells via electroporation can enhance all of the major biomass components of intact mammalian cells, we collected DNP enhancements and *T*_1_ build-up times for intact cells electroporated with increasing concentrations of AMUPol (Figure 4D and E). Relative to lysed cells, electroporated cells had lower enhancements and longer *T*_1_ values, while relative to cells incubated with AMUPol, electroporated cells had higher enhancements and shorter *T*_1_ values; DNP enhancements for proteins in electroporated cells reached a maximum value of 60 ± 6 (Figure 4D, green). However, as observed earlier, the *T*_1_ values for the AMUPol concentration (at the time of electroporation) that resulted in the highest DNP enhancements for electroporated cells were similar to those of the lysed samples (Figure 4E, green; 20 mM AMUPol, *T*_1_ = 4.0 ± 0.6 s, *n* = 4). Taken together, this indicated that the AMUPol concentration inside of electroporated cells is lower than the concentration of AMUPol that was added to the sample which reflects the difference in sample preparation. For electroporated samples, only a fraction of AMUPol enters the cells and the extracellular AMUPol is washed out of the sample after electroporation and before sample packing.

To assess the homogeneity of the AMUPol concentration throughout each biomass component in these electroporated samples, we used the regression error of the fit of the *T*_1_ data to a mono-exponential equation (Figure S4). For electroporated cells, the regression error for protein was 1.7 ± 0.4%, for nucleotide was 1.4 ± 0.4% (lower than protein, *p* = 2e-7, *n* = 12) and lipid was 1.1 ± 0.3% (lower than protein, *p* = 6e-5, *n* = 12). When compared with lysed samples, the regression error was indistinguishable for lipids (*p* = 0.38, unpaired), and the regression errors were slightly higher for protein and nucleotide (*p* < 0.05, unpaired). This indicated that the concentration distribution of AMUPol is more heterogeneous in electroporated cells than in lysed cells. When compared with the incubated but not electroporated samples, the regression errors were significantly smaller for all of the components (*p* < 0.003, unpaired). This showed that the concentration of AMUPol throughout the biomass is more homogenous in electroporated samples relative to samples that were incubated with AMUPol. Taken together, this indicates that delivery of AMUPol to mammalian cells by electroporation followed by removal of extracellular AMUPol results in polarization of all the biomass components with increased homogeneity of the AMUPol concentration throughout the sample relative to delivery of AMUPol by incubation.

### AMUPol is slowly deactivated by cellular environments at room temperature

Samples of lysed cells are flash frozen immediately upon addition of AMUPol. However, AMUPol is added to intact cells 8 minutes (for cells incubated with AMUPol) and 15 minutes (for cells electroporated with AMUPol) before starting to reduce the sample temperature at a rate of 1 °C per minute. Because sample manipulations can impact cellular integrity but exposure to cellular environments can reduce nitroxide radicals,^22^ we characterized the effect of sample preparation time on the DNP performance of AMUPol. First, we confirmed that AMUPol was not deactivated by cellular environments once samples were frozen. To do so, we measured DNP enhancements and *T*_1_ values for electroporated samples that were stored for various times after freezing and before sample measurement. These times ranged from several hours to one month. As expected, the DNP enhancements and *T*_1_ values were not sensitive to the length of time of frozen storage (data not shown). Next, to determine if AMUPol is deactivated by cellular environments at room temperature, we measured DNP enhancements and *T*_1_ values for samples prepared with different room temperature delay intervals before freezing. For lysed cells in the presence of 10 mM AMUPol, the enhancements decreased by half and the *T*_1_ values doubled after an hour at room temperature (Figure 5 and S5). When cells are electroporated in the presence of 10 mM AMUPol, allowed to recover, and then washed to remove extracellular AMUPol, the DNP enhancements decreased by more than half and *T*_1_ values more than doubled when the sample was allowed to sit for an additional 45 minutes at room temperature before freezing (Figure 5 and S5). The decreases in DNP enhancement for the lysed and electroporated cells indicate that exposure to cellular constituents at room temperature decreases the polarization ability of AMUPol and the increases in *T*_1_ values suggest that this deactivation process is via destruction of the radical. In contrast, when intact cells were incubated with 10 mM AMUPol, the enhancements and *T*_1_ values were unchanged with increasing room-temperature incubation periods. Interestingly, we know that cellular constituents deactivate AMUPol. Yet, there was no net change in DNP enhancement or *T*_1_ value with increasing incubation times in these samples. This observation could be explained if AMUPol both continuously enters and is also continuously reduced by the cell during the room temperature incubation time. Taken together, the effect of room temperature incubation times DNP enhancements demonstrates that a short time interval between exposure to AMUPol and freezing results in the best DNP performance. However, the rate of DNP performance loss was slow relative to most sample manipulation steps. Therefore, sample preparation times after addition of AMUPol should be minimized, but not at the expense of sample handling steps that can compromise sample integrity.

**Figure 5:**
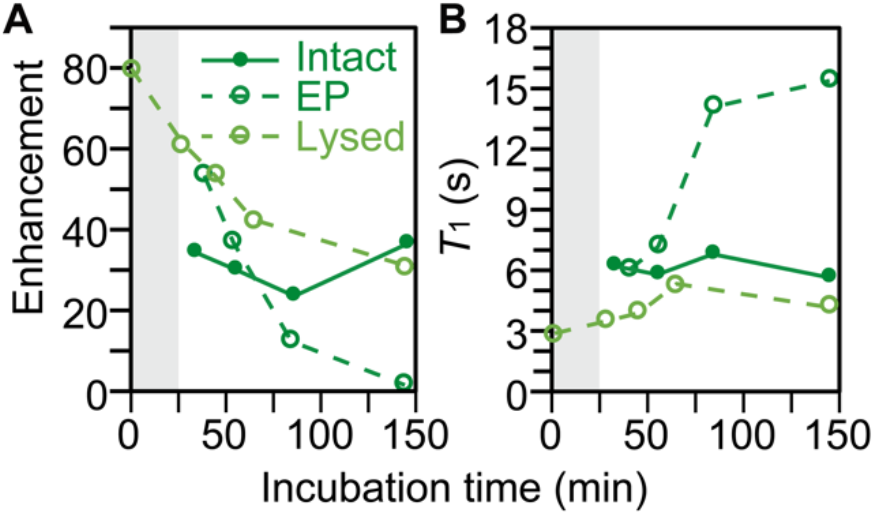
AMUPol is deactivated by cellular biomass. (A) DNP enhancements decrease and (B) *T*1 values from saturation recovery experiments increase with longer room temperature exposure to AMUPol to cellular biomass before freezing. Lysed cells with 10 mM AMUPol (light green open circles) were flash frozen after indicated room temperature incubation times. Intact cells where the 10 mM AMUPol was introduced by incubation (dark green closed circles) and where the 10 mM AMUPol was introduced by electroporation (dark green open circles) require room temperature manipulation before reducing the temperature at a rate of 1 °C/min. An additional 25 minutes (gray bar) was added to the room temperature incubation time for intact cells to visually represent this difference in sample preparation.

### AMUPol transverses the nuclear envelope of intact cells

While RNA is largely localized to the cytoplasm, DNA is mostly localized to the nucleus. RNA and DNA can be distinguished using ^13^C-^15^N correlation spectroscopy. The ^13^C chemical shift of the C_1_ carbon of the ribose in RNA molecules is 8 ppm greater than that of the C_1_ carbon of the deoxyribose in DNA molecules and the ^15^N chemical shift of the N_9_ nitrogen of the pyrimidine is ∼20 ppm greater than that of the N_1_ nitrogen of the purine (Figure 6). Thus, to determine if AMUPol can enter membrane bound organelles, we collected DNP-enhanced 2D ^13^C-^15^N correlation spectra (TEDOR)^43^ on lysed and intact HEK293 cells that were either incubated or electroporated with AMUPol and examined the deoxyribose-purine and pyrimidine regions. We observed strong DNA signals in all the samples (Figure 6), indicating that AMUPol was able to enter the nucleus in intact viable cells.

**Figure 6:**
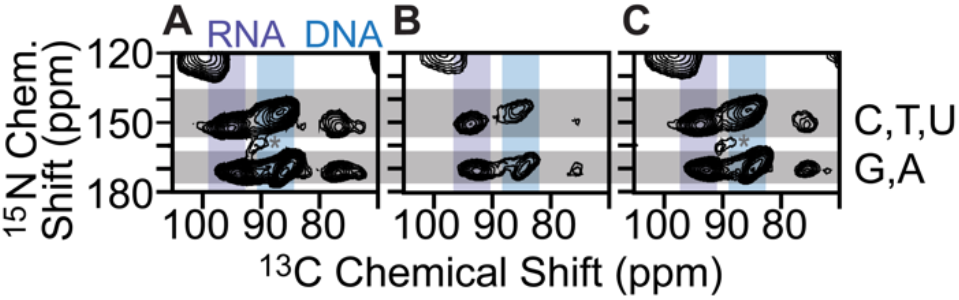
AMUPol freely enters the nucleus. Signals from RNA (purple bar) and DNA (blue bar) are resolved in the 2D TEDOR spectra A) lysed cells with 10 mM AMUPol, B) intact cells incubated with 30 mM AMUPol and C) intact cells electroporated in the presence of 20 mM AMUPol. Chemical shifts for pyrimidines and purines are indicated by the upper and lower gray bars, respectively. Spinning side bands are annotated by *.

To further distinguish the cytosolic and nucleic locations of AMUPol, we compared the normalized peak intensities of the cytosolic and nucleic components, for the lysed sample to intact samples that were either incubated or electroporated with AMUPol. The TEDOR peak intensities were normalized to either the ribose-purine peak of RNA or DNA for each sample to control for differences in DNP-enhancements and cross-polarization efficiencies. TEDOR spectra have distinct peaks for DNA, RNA, protein backbone sites, protein side chain moieties, and free amino acids (Figure S6). DNA is located only in the nucleus, while RNA, proteins and free amino acids are entirely or largely cytoplasmically localized (i.e. more than 80% of the protein content of a cell is non-nuclearly localized).^44^ In addition to the RNA and DNA ribose purine and pyrimidine peaks, we determined peak intensities for the amide-carbonyl and amide- C_α_ sites for both proteins and free amino acids as well as the carbon-nitrogen bonds in the protein side chains of arginine and glycine (Table S1). When the intensity of the amino acids peaks are compared to the ribose-purine peak of RNA all samples had similar ratios (*p* > 0.08). This indicated that the cytoplasmic distribution of AMUPol is similar for all of these samples. When the intensity of the amino acid and RNA peaks are compared to the deoxyribose-purine peak of DNA, we found that lysed and electroporated samples have indistinguishable peak intensity ratios (*p* = 0.22). However, the DNA peaks for intact cells incubated with AMUPol were less intense than expected; the ratios of peak intensities for cytoplasmic to nuclear components were larger by 60% ± 50% (*p* < 0.05); (Table S1). This suggested that the AMUPol concentration in the nucleus of cells incubated with AMUPol is lower than the concentration of AMUPol in the cytoplasm. Because DNP enhancements for cells incubated with AMUPol are independent of room temperature incubation times despite the time-dependent reduction of AMUPol by cellular biomass and these samples have heterogenous AMUPol distribution, this suggests that there is an AMUPol concentration gradient inside intact cells incubated with AMUPol. Despite the heterogeneity of the AMUPol distribution for incubated cells, however, AMUPol is able to enter the nucleus. Taken together, examination of the peak intensity ratios for the different biomass components indicates that AMUPol is able to transverse the nuclear membrane of intact cells and effectively polarizes biomolecules localized in both the cytoplasm and the nucleus.

### Sensitivity enhancements from DNP enable detection of a minority component in experimentally tractable acquisition times

To explore the sensitivity of DNP MAS NMR, we examined the signal to noise ratio (SNR) for different biomass peaks from DNP-enhanced 2D heteronuclear (TEDOR) and homonuclear (DARR) correlation spectra for lysed and intact cells that had been either incubated or electroporated with AMUPol. While RNA and DNA together comprise ∼6% of the cellular biomass, DNA alone makes up a fifth of the total nucleotide content. DNA comprises ∼1.3% of the biomass of HEK293 cells, half of which is pyrimidines.^42^ When cells were prepared without AMUPol, the peak that corresponds to the deoxyribose-pyrimidine site in DNA had an SNR of 5 after 18 hours of signal averaging. In contrast, after only two hours of signal averaging, we found that this peak for lysed HEK cells with 10 mM AMUPol had an SNR of 200, while that of intact cells incubated with 30 mM AMUPol had an SNR of 40 and that of intact cells electroporated with 20 mM AMUPol had an SNR of 110 (Figure 6). In 2D homonuclear correlation spectra, without DNP, the peak for the deoxyribose C_4_-C_5_ site had an SNR of 10 after 25 hours of signal averaging. In contrast, after only 8 hours of signal averaging, we found that this peak for lysed HEK cells with 10 mM AMUPol had an SNR of 27, while that of intact cells incubated with 30 mM AMUPol had an SNR of 18 and that of intact cells electroporated with 20 mM AMUPol had an SNR of 24. As expected, the SNRs were larger for samples with higher DNP enhancements. Comparison of the SNR of these two-dimensional spectra collected without DNP to those collect with DNP reveal a time savings of three to five orders of magnitude.

Overall, DNP-enhanced MAS NMR enabled detection of a component by both 2D homo and heteronuclear spectroscopy that comprises under 0.6% of the cellular biomass in a few hours with high signal to noise.

## DISCUSSION

NMR has the resolution and specificity to determine atomic-level protein structures of isotopically-labeled proteins in complex environments^7^ and, with the sensitivity gains conferred by dynamic nuclear polarization (DNP), NMR has the sensitivity to detect proteins at their endogenous concentrations^1^. However, just as for other structural biology techniques, DNP sensitivity enhancements are critically dependent on experimental conditions and sample composition.^35^ While some of these conditions are at least theoretically compatible with cellular viability, the effects of others on cellular sample integrity, like magic angle spinning and exposure to polarization agents, are unknown. Uncertainty about the integrity of cellular samples complicates interpretation of the data, limiting the utility of that information. We establish that sample conditions that favor efficient DNP enhancements are compatible with cellular viability. Using several measures of sample viability, we find that the rate of temperature change during freezing has the largest effect. With this in mind, we established methods that maintain cellular viability throughout the DNP NMR experiments. This allows for sample validation both before and after structural analysis. We also find that the polarization agent, AMUPol, does not affect cellular viability. Using electroporation to deliver AMUPol to cells, we find that AMUPol is distributed homogenously throughout the cytoplasm and the membrane-bound nucleus. Finally, we find that the magnitude of the sensitivity enhancements is high enough to enable detection of a protein at micromolar concentrations in experimentally tractable experimental times. We establish an approach for DNP NMR on viable cells that can be phenotyped before and after experiments to eliminate uncertainty about sample integrity.

The approach to sample preparation described in this work is likely to be easily generalized to other cultured mammalian cell lines. The freezing rate had the largest influence on post-experiment sample viability. Indeed, while protocols for cryopreservation of mammalian cells differ in the composition of the freezing media^45^, most use a standard freezing rate of 1 °C per minute^41^. Here, we eliminated carbon and nitrogen-containing components of the freezing media because doing so will facilitate spectral interpretation for samples where the target molecule is at concentrations that are low enough that signals from natural abundance components make a significant contribution to the spectra.^27^ However, DNP efficiency is unlikely to be compromised by nutrient-rich components of the freezing media since cell lysates and other complex mixtures of biomolecules support efficient DNP.^1,27,37^ Thus, as long as samples are frozen slowly and cryogenically transferred into a pre-cooled spectrometer^40^, the freezing media can be tailored to the cell line.

The method used to introduce the polarization agent affects the experimental output and therefore must be chosen to address the structural question under consideration. In-cell structural biology enables the study of protein conformation in environments that maintain the identity, stoichiometry, concentrations and organization of the myriad of biomolecules that can interact with a protein of interest.^1,7,12,18^ NMR is uniquely suited to study proteins in these complicated contexts with atomic level resolution. However, choice of approach and interpretation of the results requires consideration of the potential sources of bias. Solution state NMR experiments are biased towards the observation of mobile components, although interactions that transiently modify rotational correlation times can be inferred.^12^ MAS NMR of frozen samples are not biased by rotational correlation times but lack experimental sensitivity necessary to observe molecules at their endogenous concentrations. The sensitivity enhancements from DNP rely upon proximity to a polarization agent.^27^ Thus, DNP-enhanced MAS NMR experiments are biased towards observation of molecules that are accessible to polarization agents. In this work, we characterized samples prepared in three different ways; intact cells where AMUPol was introduced by incubation, lysed cells and intact cells where AMUPol was introduced by electroporation. For cells incubated with AMUPol, a minority of the AMUPol enters the cell; the peak intensities for DNA are lower than expected and the *T*_1_ fits indicate that the AMUPol concentration is heterogenous. While the identity, stoichiometry, concentrations and organization of the cellular components for cells incubated with AMUPol are all maintained, the experimental read-out is spatially biased. Thus, data from experiments performed on intact cells incubated with AMUPol are qualitative rather than quantitative. While any observed conformation inside cells incubated with AMUPol exists, the relative populations of different conformations cannot be inferred from peak intensities. Lysing cells results in a homogenous distribution of the polarization agent throughout the sample. For lysed cells, the identity and stoichiometry of the sample components are maintained, however lysing results in a small decrease in concentration of cellular components and loss of organization. Because the distribution of AMUPol is homogenous, the data from experiments on lysed cells are quantitative but the decrease of the concentrations and loss of organization of the environment upon lysis may alter the conformational ensemble. Introduction of the polarization agent into cells by electroporation may best address questions that require quantitative information about the structural ensemble for proteins that are potentially sensitive to not only stoichiometry and identity but also to the concentration and organization of the cellular components. Introduction of AMUPol via electroporation results in a homogenous distribution of the polarization agent throughout the cell. Using the electroporation approach for AMUPol delivery, the cellular organization is maintained and cellular propagation is unaltered. While transient permeabilization of the plasma membrane could potentially alter the identity, stoichiometry and/or concentration of cellular components, phenotyping before and after experiments eliminates uncertainty about sample integrity. Thus, structural information from experiments on electroporated cells will report quantitatively about the ensemble with minimal or no perturbation to the identity, concentration, stoichiometry or organization of the cellular constituents. Choice of methodology to introduce polarization agents therefore depends upon the structural question to be addressed.

DNP efficiencies for intact, viable samples are sufficient to collect data for proteins at their endogenous levels inside cells. Robust protocols describe successful delivery of endogenous concentrations isotopically enriched proteins into mammalian cells via electroporation.^6,12,21^ Although in cases where this technique is not suited, there are other delivery options.^8,17,46^ In this work, because DNA is spectrally resolved from other biomass components, we could specifically detect DNA, a component that comprises less than 1% of the biomass, with high signal to noise ratios in short acquisition time. The endogenous concentration of most proteins is two to three orders of magnitude lower than DNA. However, using DNA as a benchmark suggests that a protein at a micromolar concentrations could be detected by two-dimensional spectroscopy with a SNR of ∼10 in experimental acquisition times of a few days to a week. These experimental times and SNR are similar to prior reports of proteins at micromolar concentrations in complicated biological mixtures using DNP-enhanced MAS NMR.^1,21,27^ Thus, the magnitude of the DNP sensitivity enhancements in preparations of intact viable cells is sufficient to enable detection of a protein at micromolar concentrations in experimentally tractable experimental times.

A concern that plagues structural biology is that the manipulations that permit structural characterization may also alter molecular structure. Such alternations are of particular concern in cases for proteins whose conformations are environmentally sensitive, like those that can adopt more than one stable structure or contain regions that are intrinsically disordered in some settings. If cells survive structural characterization, then biological phenotypes can be assessed both before and after DNP MAS NMR. If the manipulations do not affect phenotype, then the observed structural ensembles will report on the biologically relevant conformations. Here we describe a generalizable approach for in-cell structural biology that allows both pre- and post-experimental sample validation. Such studies could significantly expand our understanding of protein conformation changes that influence cellular fate.

## Acknowledgements

This work was supported by grants from the National Institute of Health (NS-111236), the Welch Foundation (I-1923_20170325, the Lupe Murchison Foundation and the Kinship Foundation (Searle Scholars Program) to K.K.F. The authors would like to acknowledge the assistance of the UT Southwestern Live Cell Imaging Facility, a Shared Resource of the Harold C. Simmons Cancer Center, supported in part by an NCI Cancer Center Support Grant, 1P30 CA142543-01

## Materials and Methods

### Sample preparation

Human embryonic kidney 293 (HEK293) cells were cultured in ^13^C, ^15^N labelled media (BioExpress 6000 Mammalian U-^13^C, 98%; U-^15^N, 98%, Cambridge Isotope Laboratories, USA) with 10% (v/v) fetal bovine serum (FBS, qualified, Gibco) and 1% (v/v) PenStrep (Gibco) at 37 °C and 5% CO_2._ Confluent plates were harvested using Tryp-LE Express (Gibco) and BioExpress 6000 media, transferred to 15 mL conical tube and centrifuged at 233 x *g* for 5 min at 22 °C using a swinging bucket rotor (Beckman Coulter). Pelleted cells were resuspended and washed once with 1x PBS (-CaCl_2_, -MgCl_2_, pH 7.4, Gibco). AMUPol was delivered to cells in two ways: incubation and electroporation. For delivery by incubation, a 50 µL cell pellet was mixed with 50 µL perdeuterated 1x PBS (85% D_2_O + 10% H_2_O, pH 7.4) containing AMUPol (Cortecnet, USA) and 18 µL of *d*_*8*_-glycerol. The 118 µL cell suspension had a final composition of 15% (v/v) *d*_*8*_-glycerol, 75% (v/v) D_2_O and 10% (v/v) H_2_O. For delivery by electroporation, a 50 µL cell pellet was mixed with 100 µL electroporation buffer (SF cell line solution, Lonza) containing AMUPol and electroporated (HEK293 pulse sequence, Lonza 4D-nucleofactor) using manufacturer’s instructions. Post electroporation, cells were allowed to recover for 10 minutes in electroporation buffer containing AMUPol inside the tissue culture hood. Next, cells were washed twice with 50-100 µL (depending on cell pellet volume) of 1x PBS and the 50 µL cell pellet was resuspended in perdeuterated 1x PBS and *d*_*8*_-glycerol for a final composition of 15% (v/v) *d*_*8*_-glycerol, 75% (v/v) D_2_O and 10% (v/v) H_2_O.

In case of intact cells, after delivery of AMUPol, cells were transferred into 3.2 mm sapphire rotor (Bruker) by centrifugation in a swinging bucket rotor at 100 x *g* for 2 minutes at 22 °C. The supernatant was removed, and rotors were frozen at a controlled rate (–1°C/min) in ‘Cool Cell LX’ (Corning) in the –80 °C freezer for 12-16 hours. Finally, frozen rotors were transferred to liquid nitrogen storage until measurement by DNP NMR. In case of lysates, after AMUPol delivery by incubation, cells were transferred into 3.2 mm sapphire rotor (Bruker) by centrifugation in a swinging bucket rotor at 100 x *g* for 2 minutes at 22 °C. The supernatant was removed, and rotors were flash frozen in liquid nitrogen and stored till DNP NMR. During sample insert, rotors were taken out of liquid nitrogen, thawed to room temperature and then inserted into pre-cooled spectrometer at 100 K (adding one additional freeze thaw cycle to the sample handling protocol).

### Trypan Blue exclusion assay

Pelleted cells (10 µL) were diluted into 100 µL unlabeled DMEM and 10 µL of this cell suspension were mixed with 10 µL of Trypan Blue (0.4% solution). 10 µl of the Trypan Blue cell suspension was loaded onto Countess Chamber. Trypan blue membrane permeability was assessed using Countess automated cell counter (Life technologies) using the manufacturer’s protocol.

### Growth assay

Equal number of cells (0.1 million cells) were plated in 10 cm dish containing complete media (DMEM) and grown for 9-14 days (as indicated before). After cells have settled down (post 8-10 hours), media was removed to get rid of floating dead cells. 10-12 mL of DMEM is added to the 10 cm culture dish and cell growth is monitored using inverted light microscope till 100% confluency. Fitting of sigmoidal curves was performed with an equation of, where y(*t*) denotes the cell culture time *t*, a and *k* are fitting parameters, and *t*_0_ defines a lag time of *t*_L_ as *t*_L_ = *t*_0_ -2/*k*.^47^ The error range for the fitting was estimated at the 95% confidence level.

### Microscopy

HEK293 cells were grown to confluency in 10 cm dish and harvested. Cells were washed with 1x PBS twice and centrifuged at 233 x *g* for 5 minutes. 50 µL of cell pellet was resuspended in 50 µL of 1x perdeuterated PBS containing 10mM AMUPol with 15% (v/v) *d*_*8*_-glycerol. Next, the cells were centrifuged as mentioned above and the supernatant was discarded. The cell pellet was then either prepared for imaging or slow frozen (1 °C/min) and then prepared for imaging.

### Before freezing sample

Cell pellet was mixed with warm complete media (DMEM with 10% FBS and 1% Pen-Strep), centrifuged (233 x *g*, 5 min) and supernatant was discarded.

### After freezing samples

Frozen cell pellets were thawed for 2 min in a 37 °C water bath, resuspended in warm complete media (DMEM with 10% FBS and 1% Pen-Strep), centrifuged (233 x *g*, 5 min) and supernatant was discarded. The cell pellets were resuspended in 250 µL of PBS containing 4% paraformaldehyde for 30 min to fix the cells. Excess paraformaldehyde was quenched with 0.1 M glycine in PBS for 5 min. 300 µL of 0.1% Triton X-100 (in PBS) was added to the cell pellet and resuspended for 3-5 min to increase permeability followed by 1x PBS wash (3 times). Phalloidin solution [1x Phalloidin (ab176753, Abcam) + 1% BSA in PBS] with Hoechst 33342 (Invitrogen, R37605) was added to the cells and they were incubated at room temperature for 60 minutes. Cells were pelleted and rinsed 2-3 times with 1x PBS, 5 min per wash. Cell pellets were resuspended in 100 µL of PBS. 5 µL of the resuspension solution was dropped on glass slide and imaged using confocal microscopy (Zeiss LSM780 inverted). Images were analyzed with Zen Black and Fiji software (NIH).

### Cold insert

Rotors were transferred in liquid nitrogen directly into the NMR probe that was pre-equilibrated at 100 K^40^.

### DNP NMR spectroscopy

All dynamic nuclear polarization magic angle spinning nuclear magnetic resonance (DNP MAS NMR) experiments were performed on a 600 MHz Bruker Ascend DNP NMR spectrometer/7.2 T Cryogen-free gyrotron magnet (Bruker), equipped with a ^1^H, ^13^C, ^15^N triple-resonance, 3.2 mm low temperature (LT) DNP MAS NMR Bruker probe (600 MHz). The sample temperature was 104 K and the MAS frequency was 12 kHz. The DNP enhancement for the instrumentation set-up for a standard sample of 1.5 mg of uniformly ^13^C, ^15^N labeled proline (Isotech) suspended in 25 mg of 60:30:10 *d*_*8*_-glycerol:D_2_O:H_2_O containing 10 mM AMUPol was between 130 and 140. For ^13^C cross-polarization (CP) MAS experiments, the ^13^C radio frequency (RF) amplitude was linearly swept from 75 kHz to 37.5 kHz with an average of 56.25 kHz. ^1^H RF amplitude was 68∼72 kHz for CP, 83 kHz for 90 degree pulse, and 85 kHz for ^1^H TPPM decoupling with phase alternation of ± 15° during acquisition of ^13^C signal. The DNP enhancements were determined by comparing 1D ^13^C CP spectra collected with and without microwaves irradiation. For *T*_1_ measurements, recycle delays ranged from 0.1 s to 300 s. To determine the *T*_1_, the dependence of the recycle delay on both ^13^C peak intensity or volume was fit to the equation.

^13^C-^13^C 2D correlations were measured using 20 ms DARR mixing with the ^1^H amplitude at the MAS frequency. A total of 280 complex *t*_1_ points with increment of 25 μs were recorded. For ^13^C-^15^N 1D and 2D correlations, a 2 ms TEDOR sequence was applied with ^13^C and ^15^N pulse trains at 55.5 kHz and 41.7 kHz, respectively. A total of 64 complex *t*_1_ points with an increment of 80 μs were recorded. The recycle delay was 3.9 s and the same ^1^H decoupling was applied.

### DNP NMR Data analysis

For 1D experiments, the data were processed using NMRPipe^48^. The real part of the processed spectrum was exported using pipe2txt.tcl command. Peaks were integrated, and the time constants were obtained by least-squares fitting with a single-exponential function. To obtain the error estimates for DNP enhancements, the DNP enhancements were measured on three independent sample preparations for each time point. The reported errors were calculated by pooling variances across all incubation time points. DNP enhancements were determined by peak intensity. For 2D experiments, the TEDOR and DARR data were both apodized with a Lorenz-to-Gauss window function with IEN of 15 Hz and GB of 75 Hz in the *t*_1_ and *t*_2_ time domains. The experiment time of TEDOR of 32 scans were 2 h and DARR of 16 scans were 5 h. The noise level and peak height from the 2D NMR spectrum was detected by the NMRDraw software for S/N estimation.

### Quantification of the biomass components

We quantified the biomass components by integration of the featured peaks described above and then converted to biomass by multiplying the molecular weight ratio of molecular to chemical group. The peak intensities from the direct polarization experiments report on the amount of carbon atoms for each biomass component. To quantify each cellular biomass, the integrations of carbon signals have to be converted into the total mass percentage for each biomass component by multiplying a normalization factor. The normalization factors used are 10.8 (protein), 11.7 (RNA), and 3.12 (lipid). Details for obtaining these factors were described as follows. The signal reporting on protein at 178 ppm arises from two carboxyl carbons atom (24 g/mol) per aspartic acid and glutamic acid, and one carboxyl carbon atom (12 g/mol) per the rest of each amino acid. The normalization factor for protein is the average of ratios of total molecular weight to the carboxyl carbon molecular weight for each amino acid. The signal reporting on RNA was integrated in the range between 134.5 ppm and 145 ppm. Two carbon atoms per UMP and CMP and three carbon atoms per AMP and GMP are located in this region. The average of each NMP molecular weight over carbon mass was used as the normalization factor for RNA. For lipid component, the carbons at 37 ppm are -CH_2_- that is mainly located in the fatty acid chain. The average carbon numbers in the fatty acid chain were calculated to be 17.3 for HEK293 cells,^49^ where the average carbon numbers at 37 ppm per fatty acid chain is 15.3. Cholesterol (4 carbons at 37 ppm), sphingolipids (average of 27.3 carbons), phosphatidylcholine (average of 30.7 carbons), and phosphatidylethanolamine (average of 30.7 carbons) are most abundant in HEK293 cells and were used to calculate the average normalization factor of lipid components. The calculation of the normalization factors for each biomass component could vary slightly, based on whether the biomass of each individual amino acids, nucleic acids, or lipids are counted in or not.^42^ With additional weighing factor, the normalization factors became 10.0 (protein), 15.0 (nucleic acid), and 3.43 (lipid). Both normalization methods show that NMR-based biomass quantification could recapitulate literature values^42^ using different biomass quantification approaches.

