## Supplemental figures and table for "In-Cell Sensitivity-Enhanced NMR of Intact Living Mammalian Cells"

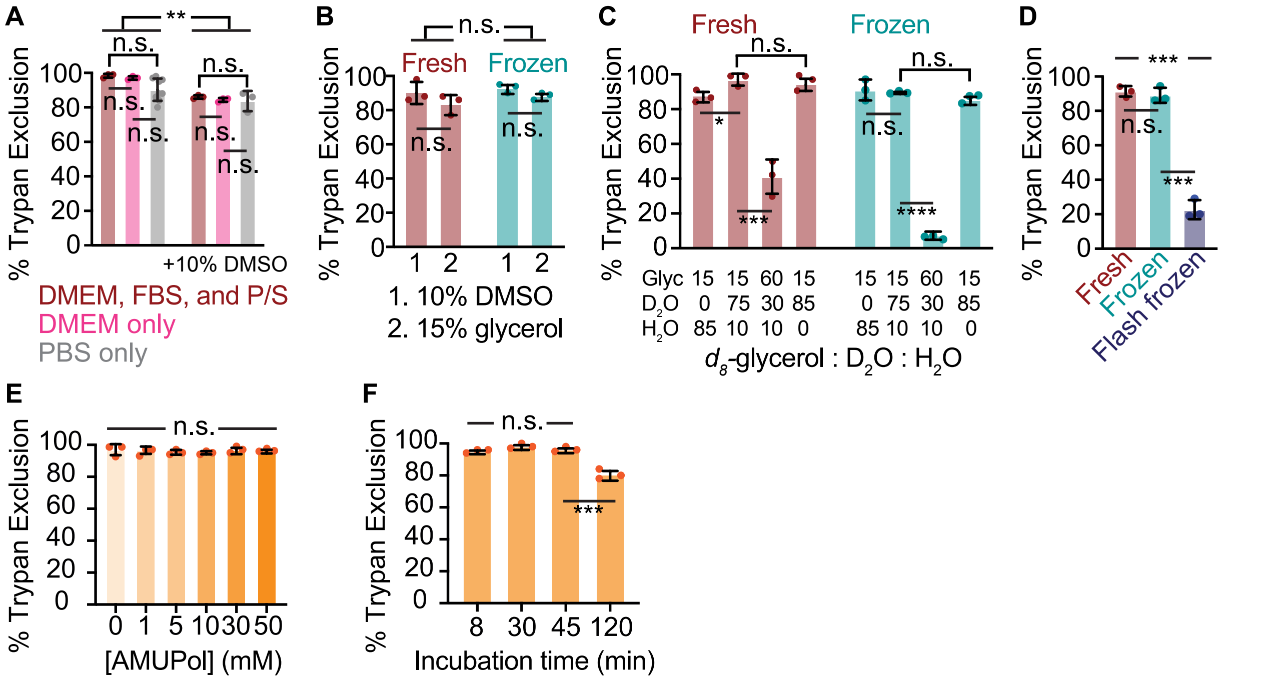


**Figure S1. HEK293 cells can survive DNP NMR process** (A) Freezing media containing only PBS with 10% DMSO can cryopreserve HEK293 cells in comparison with the complete media (DMEM with 20% FBS and 1% PenStrep). However, the freezing process decreases viability regardless of the freezing media components ($p$ = 0.004, Wilcoxon's matched pair signed rank test). (B) 15% glycerol can be used as a cryoproectant with the same efficiency as 10% DMSO. (C) Increase in deuteration percentage does not affect HEK293 cell viability. However, increasing $d_8$-glycerol percentage (v/v) from 15% to 60% reduces cell viability from 97 ± 3.5% to 41 ± 9.9% before freezing ($p$ = 0.001, Student's t-test) and 89.3 ± 0.6% to 7.7% ± 2.1% after freezing conditions ($p$ = 2e-6, Student's t-test). (D) Flash freezing of HEK293 cells significantly reduces viability (22.7 ± 5.5%) compared to controlled freezing at 1 °C/min (89 ± 4.4%) ($p$ = 1e-4, Student's t-test). (E) Increase in AMUPol concentration till 50 mM does not affect HEK293 cell viability ($p$ = 0.749, One-way ANOVA). (F) Increase in incubation time till 45 min does not affect HEK293 cell viability as well ($p$ = 0.1, One-way ANOVA). However, viability starts to decline by 120 min significantly (p=0.001, Student's t-test).

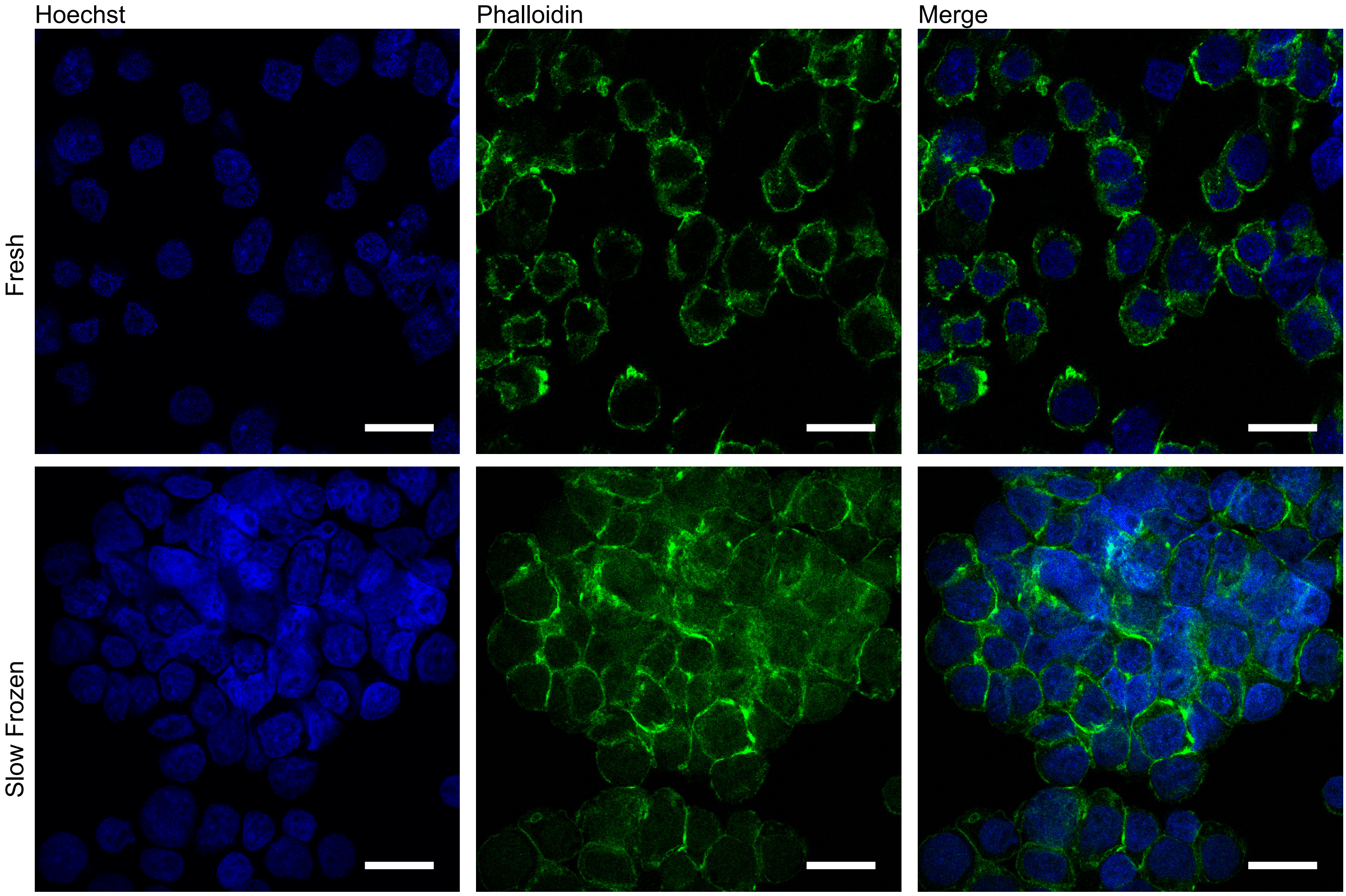


**Figure S2. Hoechst 33342-Phalloidin staining of HEK293 cell pellets.** Cells were mixed with DNP buffer and 10 mM AMUPol, fixed with 4% PFA, and stained with Hoechst and Phalloidin before (Fresh) and after freezing (Slow Frozen). The scale bar is 20 μm.

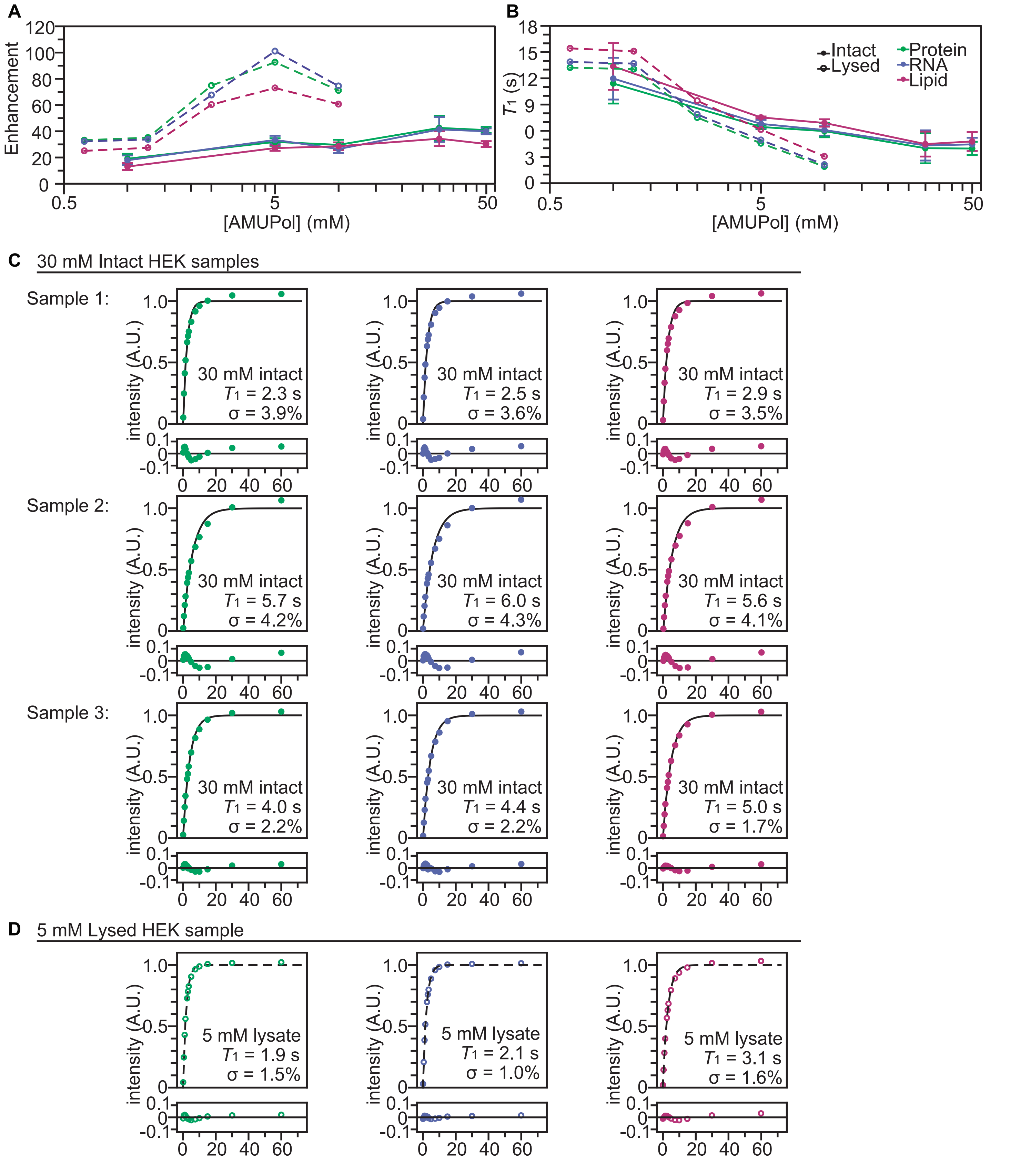


**Figure S3: AMUPol Permeabiltiy.** AMUPol concentration dependence of enhancements of protein (green), RNA (purple), and lipid (pink). DNP enhancement (A) and $T_1$ (B) of lysed HEK293 samples (dashed lines and open dots) and of live HEK293 cells (solid lines and filled dots). (C) The $T_1$ fitting results of three 30 mM AMUPle incubated HEK293 samples, and (D) the $T_1$ fitting results of the 5 mM AMUPol in lysed HEK293 sample.

20 mM Electroporated HEK sample

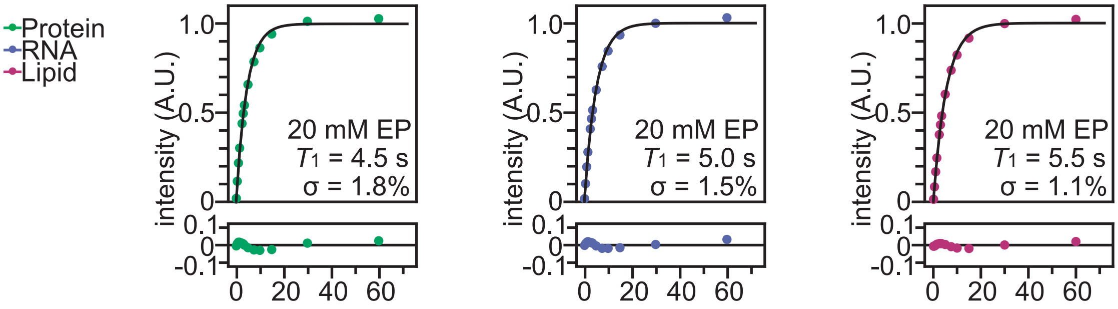


**Figure S4: AMUPol homogeneity and t1 analysis on electroporated HEK sample with 20 mM of AMUPol.** Saturation recovery analysis for $T_1$ evaluation on protein (green), RNA (purple), and lipid (pink). The enhancements on protein, RNA, and lipid are 62x, 59x, and 55x, respectively.

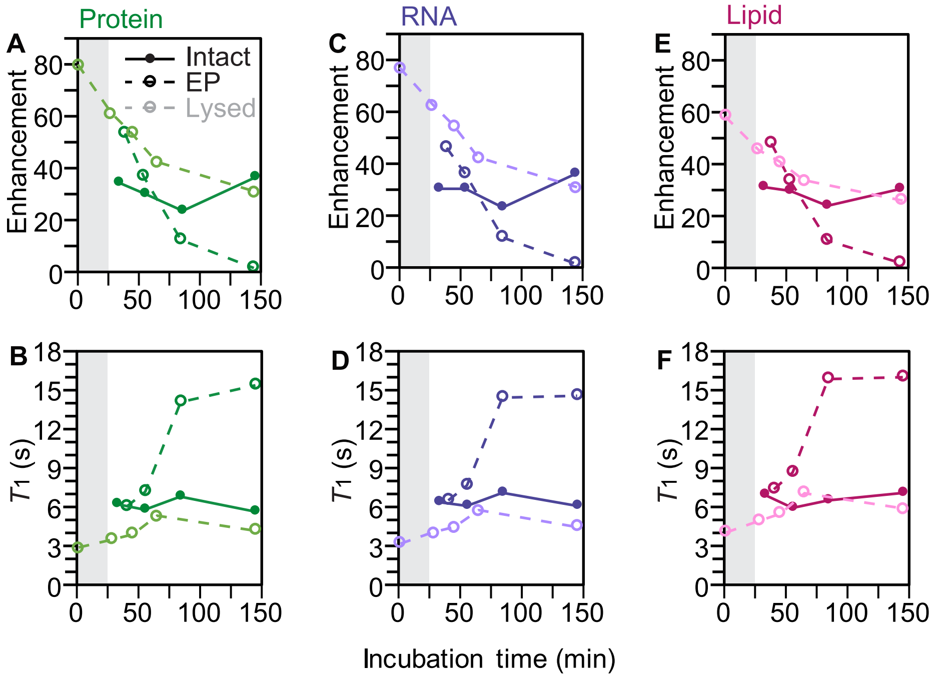


**Figure S5: AMUPol is deactivated by cellular biomass.** (A, C, E) DNP enhancements decrease and (B, D, F) $T_1$ values from saturation recovery experiments increase with longer room temperature exposure to AMUPol to cellular biomass before freezing. DNP enhancements and $T_1$ values are reported for the major biomass components; protein (A, B), RNA (C, D) and lipids (E, F). Lysed cells with 10 mM AMUPol (open circles and dashed lines) were flash frozen after indicated room temperature incubation times. Intact cells where the 10 mM AMUPol was introduced by incubation (closed circles and solid lines) and where the 10 mM AMUPol was introduced by electroporation (open circles and dashed lines) require room temperature manipulation before reducing the temperature at a rate of 1 °C/min. An additional 25 minutes (gray bar) was added to the room temperature incubation time for intact cells to visually represent this difference in sample preparation.

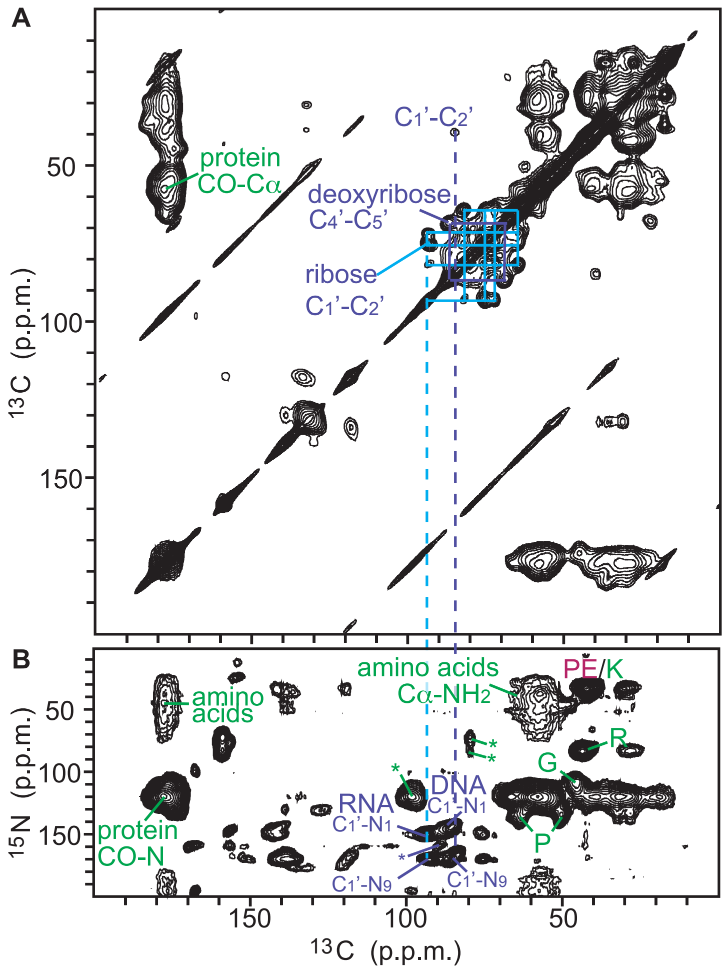


**Figure S6. 2D NMR spectrum.** Homonuclear and heteronuclear correlation spectra of uniformly labeled HEK293 cells electroporated in the presence of 20 mM AMUPol A) Select $^{13}C$-$^{13}C$ correlations from carbons in the ribose and deoxyribose rings of RNA and DNA are indicated in the $^{13}C$-$^{13}C$ DARR spectra. B) Selected $^{13}C$-$^{15}N$ correlations from protein back bone and side chains (green), nucleotides (blue) and lipids (pink) are indicated in the $^{13}C$-$^{15}N$ TEDOR specta. Signal to noise ratios for selected peaks are reported in Table S1.

### Table S1: Signal to noise ratios for biomass components in two-dimensional spectra.

| Samples | No DNP | | | Lysed, $\varepsilon$ = 81 | | | Incubation, $\varepsilon$ = 32 | | | Electroporation, $\varepsilon$ = 53 | | |
|---|---|---|---|---|---|---|---|---|---|---|---|---|
| Biomass components | $t_2$ (ppm) | $t_1$ (ppm) | SNR | $t_2$ (ppm) | $t_1$ (ppm) | SNR | $t_2$ (ppm) | $t_1$ (ppm) | SNR | $t_2$ (ppm) | $t_1$ (ppm) | SNR |
| TEDOR | NS = 288 | | | NS = 32 | | | NS = 32 | | | NS = 32 | | |
| DNA $C_1$'-$N_1$ | 86.0 | 144.5 | 5 | 85.5 | 144.4 | 199 | 85.7 | 144.8 | 39 | 85.7 | 144.9 | 109 |
| DNA $C_1$'-$N_9$ | 85.2 | 171.7 | 5 | 84.5 | 169.8 | 274 | 84.4 | 169.6 | 47 | 84.9 | 170.5 | 144 |
| RNA $C_1$'-$N_1$ | 93.2 | 152.1 | 10 | 93.8 | 151.0 | 284 | 93.7 | 150.9 | 70 | 93.9 | 151.2 | 168 |
| RNA $C_1$'-$N_9$ | 92.6 | 170.7 | 6 | 92.7 | 170.8 | 474 | 92.7 | 171.0 | 115 | 92.7 | 170.9 | 254 |
| Amino Acid $C_\alpha$-N | n. d. | n. d. | n. d. | 56.9 | 37.2 | 146 | 56.6 | 38.6 | 76 | 57.8 | 38.3 | 72 |
| Amino Acid CO-N | n. d. | n. d. | n. d. | 177.8 | 41.1 | 62 | 177.8 | 40.0 | 24 | 178.1 | 45.0 | 38 |
| Protein $C_\alpha$-N | 57.9 | 120.1 | 49 | 58.0 | 120.2 | 2357 | 58.0 | 120.3 | 680 | 58.0 | 120.4 | 1321 |
| Protein CO-N | 177.8 | 119.9 | 36 | 177.9 | 120.0 | 1876 | 177.9 | 120.1 | 586 | 177.9 | 120.1 | 1137 |
| Arg $C_\zeta$-$N_\xi$ | 43.7 | 83.4 | 14 | 43.5 | 83.3 | 593 | 43.6 | 84.1 | 160 | 43.6 | 84.1 | 320 |
| Gly $C_\alpha$-N | 45.1 | 108.2 | 11 | 45.4 | 108.9 | 350 | 45.4 | 109.5 | 100 | 45.6 | 109.0 | 197 |
| DARR | NS = 80 | | | NS = 16 | | | NS = 16 | | | NS = 16 | | |
| DNA $C_4$'-$C_5$' | 86.7 | 68.7 | 10 | 88.3 | 69.8 | 27 | 87.0 | 68.6 | 18.1 | 86.6 | 68.5 | 24 |

Direct dimension ($t_2$): $^{13}$C chemical shift in p. p. m. (DSS referenced) for both TEDOR and DARR experiments.
Indirect dimension ($t_1$): $^{15}$N chemical shift in p. p. m. for TEDOR experiments and $^{13}$C p. p. m. for DARR experiments.
Enhancement of protein ($\varepsilon$): Calculated from the CO peak intensity ratio of $^1$H-$^{13}$C 1D CP experiments ($d_1$ = 10 s) with and without microwave.
